# Budding yeast complete DNA replication after chromosome segregation begins

**DOI:** 10.1101/407957

**Authors:** Tsvetomira Ivanova, Michael Maier, Alsu Missarova, Céline Ziegler-Birling, Lucas B. Carey, Manuel Mendoza

## Abstract

To faithfully transmit genetic information, cells must replicate their entire genome before division. This is thought to be ensured by the temporal separation of replication and chromosome segregation. Here we show that in a substantial fraction of unperturbed yeast cells, DNA replication finishes during anaphase, late in mitosis. High cyclin-Cdk activity inhibits replication in metaphase, and the decrease in cyclin-Cdk activity during mitotic exit allows DNA replication to finish at difficult-to-replicate regions. Replication during late mitosis correlates with elevated mutation rates, including copy number variation. Thus, yeast cells temporally overlap replication and chromosome segregation during normal growth, possibly allowing cells to maximize population-level growth rate while simultaneously exploring greater genetic space.

**One Sentence Summary:** Completion of DNA replication is coupled to downregulation of Cyclin-Dependent Kinase during mitotic exit.

## Main text

Eukaryotic cells must complete DNA replication before chromosome segregation in order to maintain genomic stability. Complete replication is thought to be ensured by the temporal separation of DNA synthesis (S-phase) from mitosis (M-phase) (*1*). The ordering of S and M phases is established by increasing levels of cyclin-dependent kinase (Cdk) activity during the cell cycle (*2*) and is enforced by checkpoints that inhibit chromosome segregation when cells are exposed to severe replication stress (*3*). However, some yeast mutants, and cancer cells exposed to mild DNA replication stress, perform DNA synthesis in mitosis and possibly even in the subsequent G1 (*4, 5*), suggesting that DNA synthesis and mitosis may not be fully incompatible. Supporting this view, several lines of evidence suggest that budding yeast lack a checkpoint to detect if DNA replication has completed before entry into mitosis (*6*–*10*). Thus, to what extent eukaryotic cells temporally separate DNA synthesis and segregation under physiological conditions remains an open question.

To directly test if DNA synthesis occurs during mitosis in unstressed cells we arrested yeast in metaphase via depletion of the anaphase promoting complex activator Cdc20 and measured incorporation of the nucleotide analogue 5-ethynyl-2’-deoxyuridine (EdU) as cells were, or were not, released into a G1 arrest. Cells held in metaphase showed no nuclear EdU signal after a 60-minute pulse, whereas cells released from metaphase into G1 arrest incorporated EdU into the nucleus **(Fig.1A, S1A)**. EdU incorporation was higher in G1 than in metaphase cells in both DAPI-rich and DAPI-poor nuclear regions, which contained the nucleolar marker Net1 (**Fig. S1B**). Freely-cycling unstressed cells showed significant nuclear EdU incorporation in late mitosis and G1, whereas mitotic cells with actively segregating nuclei did not (**Fig. S2**). Further, 45% of log-phase cells entered anaphase with single-stranded DNA, detected as Replication Protein A (RPA) foci (Rfa2-GFP) **(Fig. S3)**. These observations suggest that metaphase is refractive to nuclear DNA synthesis, but that some DNA synthesis occurs between metaphase and the following G1 in freely cycling, unstressed cells.

Next, we evaluated whether DNA synthesis during mitosis is important for chromosome segregation and cell division. We arrested cells in metaphase by depletion of Cdc20 or by treatment with nocodazole, an inhibitor of microtubule polymerization. Upon release we inhibited DNA synthesis by treatment with the ribonucleotide reductase inhibitor hydroxyurea (HU) or by inactivating DNA replication factors with temperature-sensitive (ts) mutations, including DNA polymerase delta and epsilon, and the GINS complex component *PSF2* (**Fig. 1B, C**). In all of these cases, disruption of DNA synthesis during mitosis delayed or inhibited nuclear and cell division (visualized with the Histone H2B (Htb2)-mCherry reporter and the membrane marker GFP-CAAX, respectively) and triggered long-lived chromatin bridges (**Fig. 1B-D, S4**). Disruption of mitotic DNA synthesis also delayed cytokinesis in cells without tagged histones, although with reduced severity in response to HU **(Fig. 1C, S4E)**. Ablation of the *RAD9* checkpoint, which prevents cell cycle progression in response to DNA damage during both S phase and anaphase (*11*), abolished nuclear division delays in response to challenges in DNA synthesis during mitosis (**Fig. S4A-B**). These results suggest that mitotic DNA synthesis promotes timely chromosome segregation.

**Fig 1.**
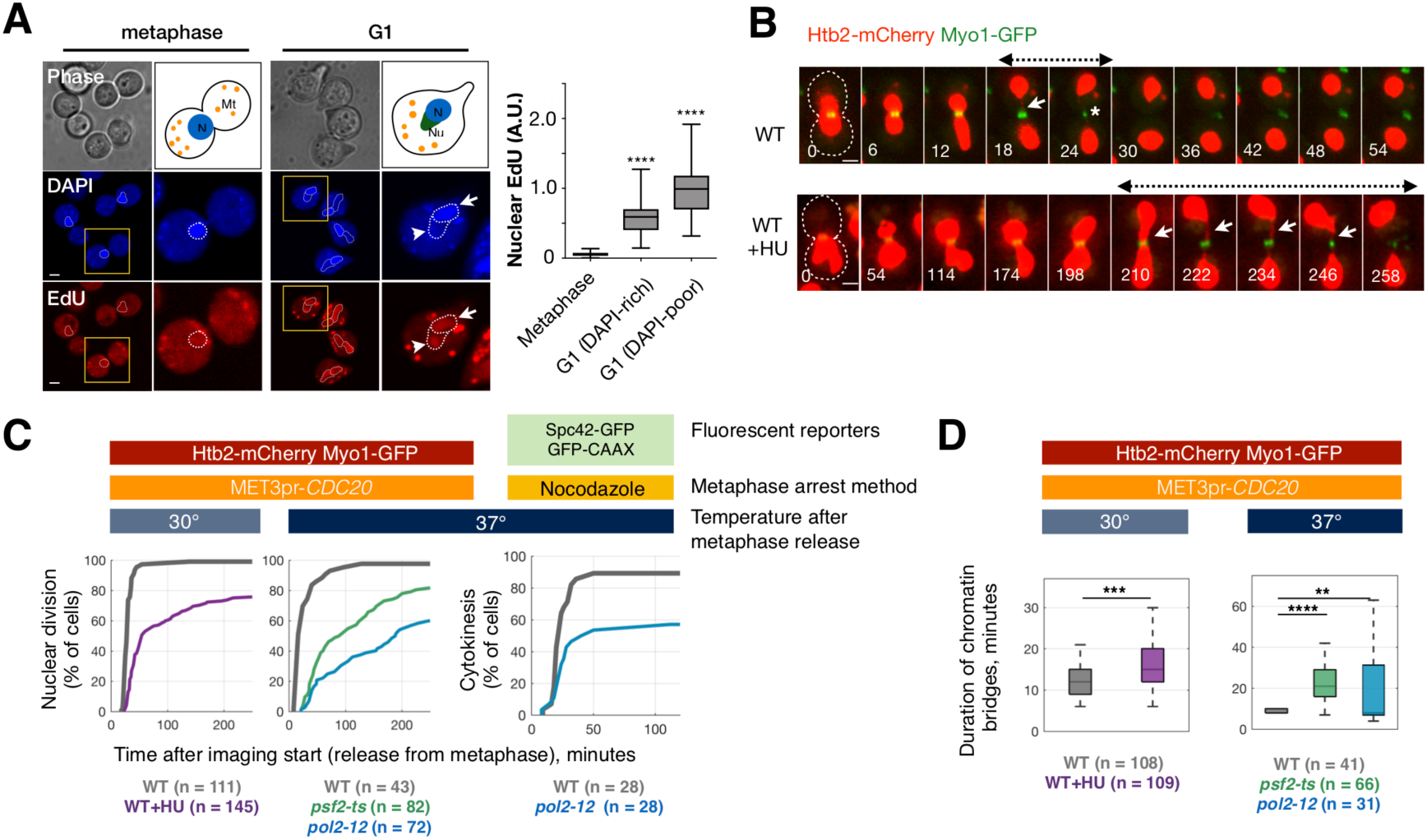
DNA synthesis in late mitosis promotes timely nuclear division. (A) *MET3pr-CDC20* cells grown for 3.5 hours in +Met medium to block them in metaphase were treated with EdU as they were either kept in metaphase or released into alpha factor-containing-Met medium for 60 minutes to arrest cells in G1. Cells blocked in G1 showed higher nuclear EdU incorporation than metaphase cells, in both DAPI-rich (arrows) and DAPI-poor regions (arrowheads). N, nucleus; Nu, nucleolus; Mt, mitochondria. ****, p<0.0001, t-test compared to metaphase. N= 27 cells / category. **(B)** *MET3pr-CDC20* cells arrested in metaphase as in (A) were released from the metaphase block in -Met medium (-HU), or treated with 100 mM HU for 30 minutes and released in HU-containing-Met medium (+HU). Arrows point to chromatin bridges labeled with Htb2-mCherry; chromatin bridge lifetime is indicated by dashed lines. Asterisks mark bridge resolution (nuclear division) and contraction of the actomyosin ring labeled with Myo1-GFP. Images were acquired every 6 minutes. The time relative to imaging start is indicated in minutes. **(C-D)** The time of nuclear division (i.e. bridge resolution) (C) and bridge lifetime (D) for cells either released from a metaphase arrest in the presence of HU at 30 °C, or arrested in metaphase at 25 °C and released at 37 °C to inactivate DNA replication. The number of cells (n, pooled from at least two independent experiments) is indicated. **** p < 0.0001; *** p < 0.001; ** p < 0.01, Student T-test. Scale bars in A, B: 2 μm.

Surprisingly, cells arrested in metaphase for prolonged periods of time did not undergo DNA synthesis during the arrest (**Fig. 1A**). We hypothesized that high Cdk activity inhibits DNA synthesis in metaphase, and that the inhibition of Cdk during mitotic exit enables synthesis to complete. To test this, we examined cells arrested in late anaphase with separated nuclei and high levels of mitotic cyclins by inactivation of the Mitotic Exit Network (MEN) (ts mutants in *TEM1, CDC15*, or *DBF2*) (*12*). MEN mutants displayed a high frequency of chromatin bridges visualized with Htb2-mCherry or the DNA dye YOYO-1, indicative of incomplete chromosome segregation in at least 40% of late anaphase cells **(Fig. 2A, S5)**. Moreover, time-lapse imaging of fluorescent loci in chromosome XII showed defects in the segregation of telomere-proximal regions in MEN mutant cells (**Fig. S6**). Chromatin bridges were stable in MEN-arrested cells (*cdc15-as1* treated with the ATP analogue 1-NA-PP1) even after inactivation of the cohesin Scc1p (**Fig. S7**). MEN reactivation (by washout of 1-NA-PP1) led to the resolution of chromatin bridges before cytokinesis even in the absence of Topoisomerase II activity (**Fig. 2B**). Thus, MEN bridges are not due to persistent cohesion or catenation between replicated sister DNA molecules. However, timely bridge resolution after MEN reactivation required DNA polymerase delta, indicating that MEN-deficient bridges require DNA synthesis for their resolution **(Fig. 2B)**. Consistent with this, RPA foci indicative of ssDNA are present in most MEN-deficient cells during anaphase **(Fig. 2C)**.

**Fig 2.**
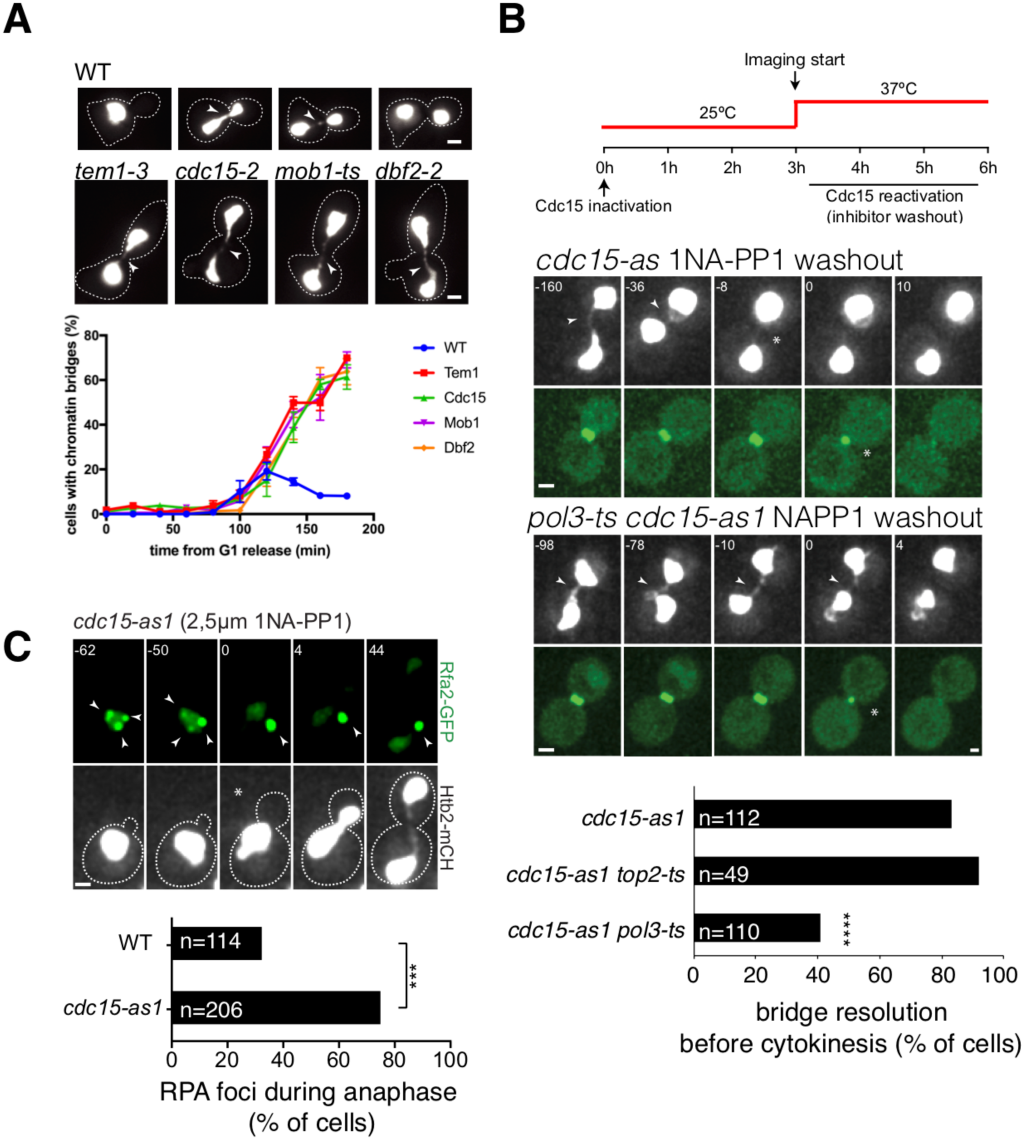
Segregation of chromatin bridges in anaphase requires the mitotic exit network and DNA synthesis. (A) Quantification of chromatin bridges in MEN mutants. Cells of the indicated strains expressing Htb2-mCherry were arrested in G1 at 25 °C, released from the block at 37 °C, fixed at the indicated times, and imaged (see *Methods*). Images depict wt cells during the time course and MEN mutant cells 2-3 hours after release from the G1 block. Arrows mark anaphase bridges. At least 100 cells were imaged per time point. The graph shows mean and SEM of 3 independent experiments. **(B)** MEN bridges require DNA synthesis but not topoisomerase II for their resolution. *Top*: experimental set-up to assess chromatin bridge resolution after MEN reactivation (see Methods). *Middle:* images of representative MEN mutants expressing Htb2-mCherry and Myo1-GFP. Arrowheads point to chromatin bridges and asterisks mark actomyosin ring contraction. Numbers indicate time (min) relative to cytokinesis. *Bottom:* Quantification of bridge resolution frequency during 3 hours following washout of 1-NA-PP1. The number of cells (n), pooled from three (*cdc15-as1 top2-4* and *cdc15-as1 pol3-ts)* or six independent experiments (*cdc15-as1*), is indicated. ****, P< 0.0001; ns, non-significant, p>0.05, Fisher’s exact test relative to *cdc15-as1*. **(C)** RPA foci persist into anaphase in MEN mutants. 1-NA-PP1 (2.5 μM) was added to mid-log phase cells (Htb2-mCherry Rfa2-GFP) at 25°C and imaged as in (B). Representative cells (WT and *cdc15-as1* mutants) are shown. Arrowheads point to Rfa2-GFP foci. The graph shows the percentage of cells that displayed RPA foci during the first 20 min after anaphase entry. The number of cells (n) pooled from three independent experiments, is indicated. ***, p < 0.0001, Fisher’s exact test. Scale bars: 2 μm.

DNA synthesis during late mitosis may reflect mitotic repair of already replicated DNA, mitotic replication of diverse genomic regions, or mitotic replication of specific genomic regions. To distinguish between these possibilities, we used Illumina sequencing to measure DNA copy number (*13*) in cells arrested in (i) G1, (ii) metaphase, via depletion of Cdc20, or (iii) late anaphase/telophase, via inactivation of MEN (*dbf2-2*). To obtain >95% synchrony we isolated mitotically arrested cells by sucrose gradient centrifugation and used the fraction with the highest synchrony for DNA extraction **(Fig. S8, Supplementary Table S1** and **Methods)**. From the copy number ratios between mitotic and G1-arrested cells we calculated the percentage of cells in which each region of the genome is under-represented during mitosis (see Methods). We excluded long repeats (rDNA and telomeres) from our analysis, as in these cases DNA copy number cannot be distinguished from repeat copy number. Nonetheless, we found that chromosome ends were under-represented in a high percentage of mitotic cells: on average, 60 kb at the end of each chromosome are under-represented in metaphase **(Fig. 3A, S9, 10A)**. Over 40% of cells have lower metaphase copy number in the 1 kb closest to each telomere **(Fig. S10B)**. Certain regions distant from telomeres were also under-represented (**Fig. 3B, 3C, Supplementary Table S2**). Regions under-represented in mitosis corresponded to a subset of late-replicating regions **(Fig. 3D, S11)**. We therefore refer to these regions as “under-replicated in mitosis”. Difficult-to-replicate regions such as those with G-quadruplexes, transposable elements and fragile sites (*14*) showed significant under-replication in metaphase (**Fig. 3E**). Under-replication was higher in metaphase (*CDC20* depletion) than in late anaphase (*dbf2* inactivation) (**Fig. 3A**). Moreover, under-replication for transposable elements and fragile sites becomes negligible in late anaphase (**Fig. S12**), suggesting that these regions complete replication in anaphase.

**Figure 3.**
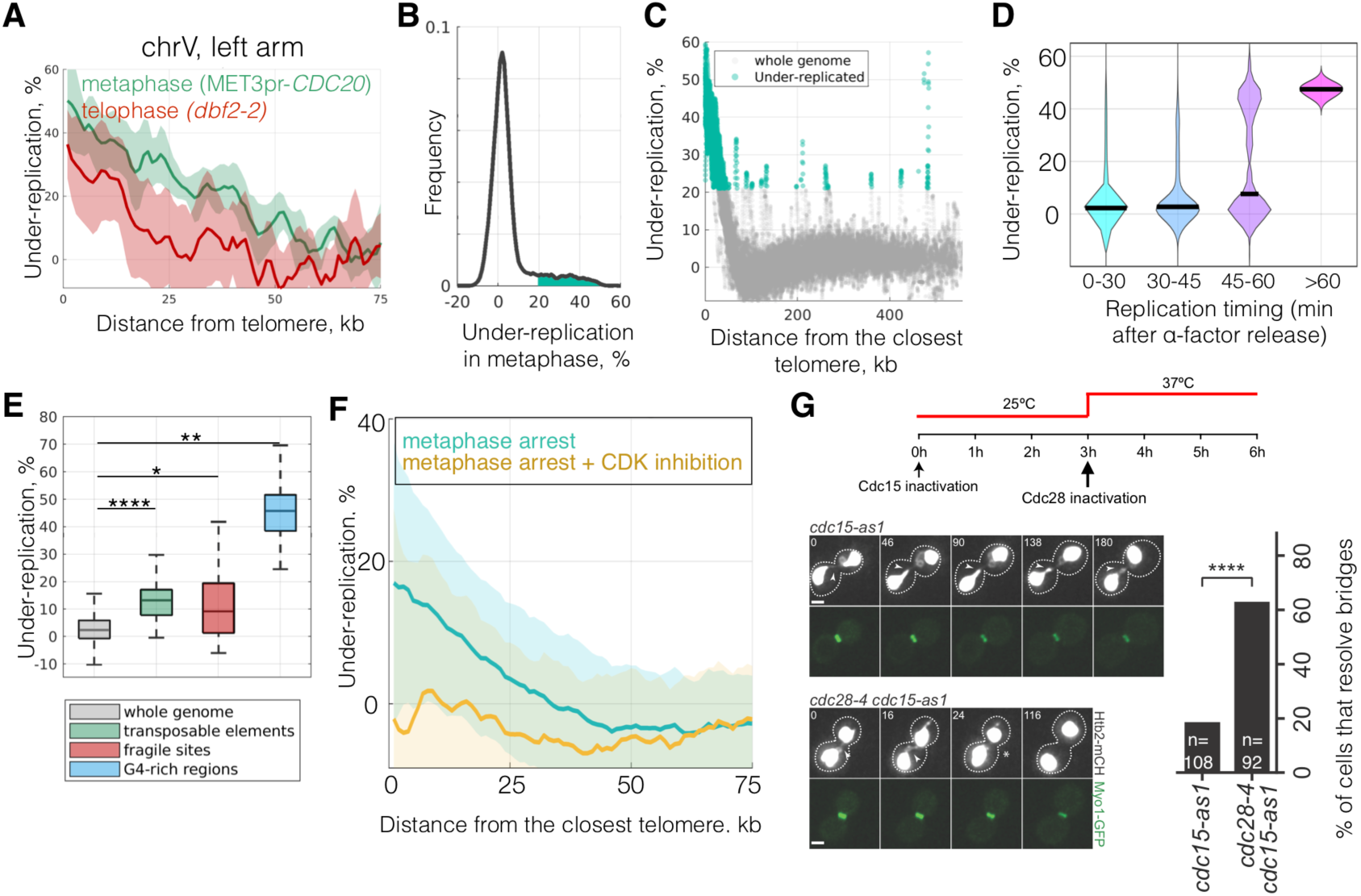
Completion of replication of subtelomeric regions and difficult-to-replicate sequences requires the drop in Cdk activity that occurs during mitotic exit. (A) Under-representation of subtelomeric regions in chromosome V for metaphase (MET3pr-*CDC20*, in green) and late anaphase (*dbf2-2*, in red) arrests. Shadows correspond to standard deviation across biological replicates (6 for MET3pr-*CDC20*, 5 for *dbf2-2*). **(B)** Distribution of sequences under-represented in metaphase across the whole genome, with values greater than a threshold of 21% (see Methods) shaded in green. **(C)** Under-representation values for all 200 bp windows throughout the genome, with significantly underrepresented genomic regions colored green. **(D)** Under-represented genomic regions in metaphase correspond to late-replicating regions. All 200-bp windows of measured under-replication split into bins based on their replication timing (data from (*23*)). **(E)** Regions with high frequency of G-quadruplexes, transposable elements and fragile sites exhibit higher under-replication in metaphase (* p < 0.05; ** p < 0.01; *** p < 0.001; **** p < 0.0001, Student T-test). **(F)** Genome sequencing was performed in metaphase-arrested (Cdc20-depleted) cells before and after inhibition of Cdk using the ATP analogue-sensitive mutant *cdc28-as1* (see Methods). **(G)** Inactivation of Cdk function allows chromatin bridge resolution in MEN-deficient (*cdc15-as* +1-NA-PP1) cells. Arrowheads mark chromatin bridges and asterisks bridge resolution. Time 0 corresponds to start of imaging (temperature shift). n=number of cells is indicated. Cells were pooled from three independent experiments. ****, p < 0.0001, Fisher’s exact test. Scale bar: 2 μm.

To directly test if high mitotic Cdk activity inhibits DNA synthesis in difficult-to-replicate genomic regions we determined DNA copy number in Cdc20-depleted cells after inactivation of Cdk using the *cdc28-as* allele. Inactivation of Cdk enabled Cdc20-depleted cells to finish replication of chromosome ends **(Fig. 3F).** Furthermore, Cdk inactivation (*cdc28-4*) also enabled chromatin bridge resolution independently of cytokinesis in MEN-deficient cells **(Fig. 3G)**. Consistent with this, over-expression of the mitotic cyclin *CLB2* in S phase slowed completion of DNA replication **(Fig. S13)**. Therefore, Cdk inactivation is required to complete replication of specific regions during mitotic exit, preventing the formation of stable chromatin bridges.

There is some correlation between replication timing and mutation rates in yeast and human cells (*18, 19*) and a strong relationship between distance to the telomere and mutation rate in budding yeast (*20*). Deletion of genes that are replicated in late mitosis has no apparent fitness cost, consistent with these genes being dispensable under non-challenging conditions (**Fig. 4A**). Interestingly, genes in regions replicated during mitosis exhibited higher levels of intraspecies genetic diversity at the single-nucleotide level, for small insertions and deletions (<50 nt), and at the level of gene gain and loss **(Fig. 4B)**. Further, gene ontology analysis shows that genes under-replicated in mitosis are mostly subtelomeric, and are enriched in carbon signaling and transport functions (**Fig. 4C** and **Supplementary Table S3**). Thus, replication during late mitosis may result in increased mutation rates of specific classes of genes.

**Fig 4:**
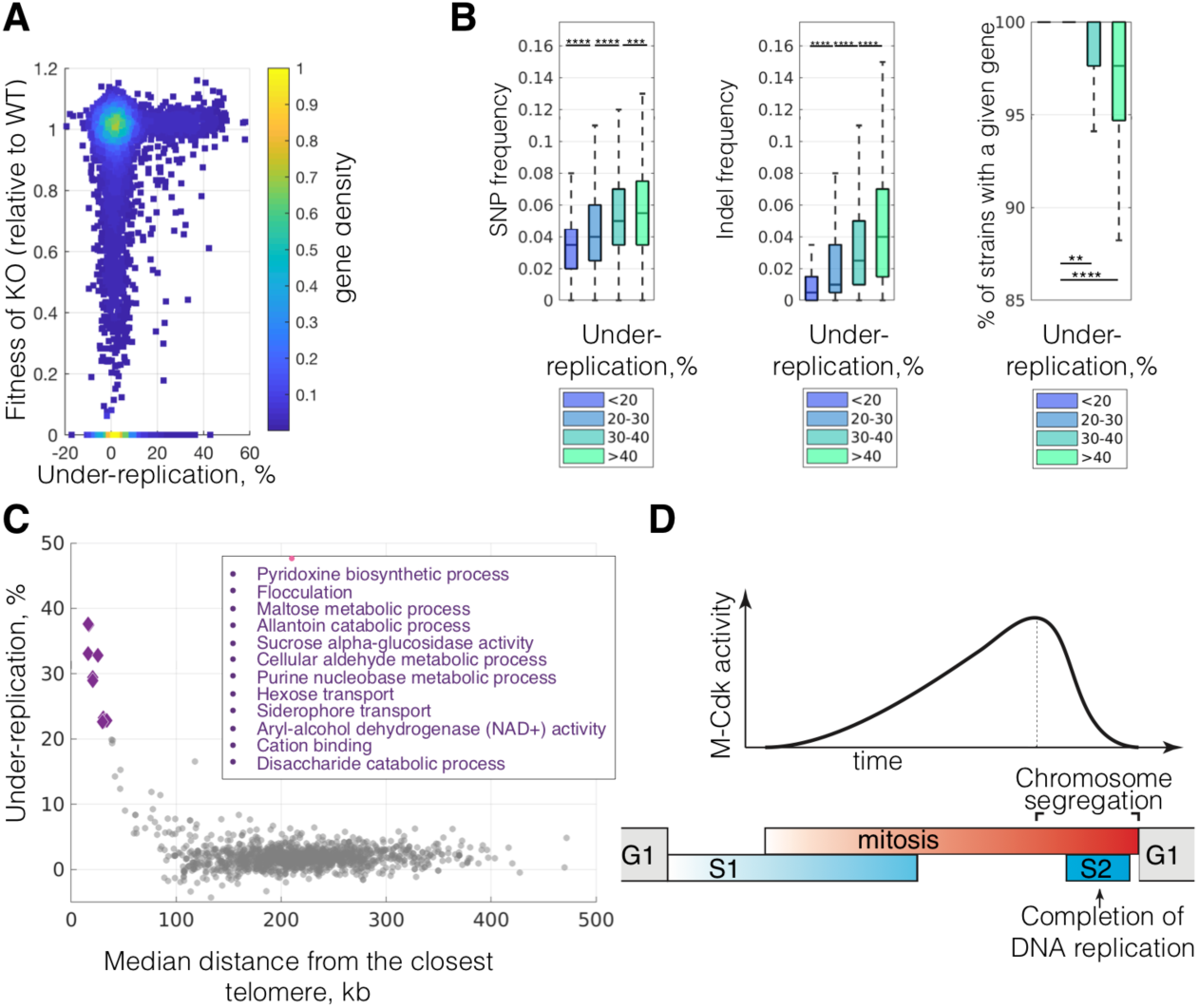
Regions replicated after metaphase are non-essential and have high rates of evolutionary divergence. (A) Each point represents one gene, showing the % under-replication in metaphase-arrested cells and the fitness of the deletion mutant (0 for essential genes). **(B)** Regions that are under-replicated in metaphase have higher SNP and InDel frequencies, and are more likely to be lost, across 85 *S. cerevisiae* strains (data from (*24*)) (** p < 0.01, **** p < 0.0001, Student T-test). **(C)** Each point is the mean value for under-replication and distance to the nearest telomere for all genes contained in each gene ontology (GO) term. Analysis in (A-C) was performed using data from metaphase arrested *MET3pr-CDC20* cells. **(D)** Budding yeast complete DNA replication after chromosome segregation begins. The order of cell cycle phases is represented by grey (G1), blue (S1, S2) and red (M) bars. S phase is divided into pre-mitotic S1, in which the bulk of DNA replication occurs, and S2, which overlaps with chromosome segregation and in which subtelomere, transposons and fragile sites are replicated in a fraction of cells. DNA replication is inhibited by high M-Cdk levels during metaphase.

Together, these results suggest that a substantial fraction of wild-type unstressed cells finish replication of specific chromosome regions late in mitosis, long after the initiation of chromosome segregation (**Fig. 4D**). We speculate that replication forks likely stall at difficult-to-replicate sequences, such as G-quadruplexes, and that their replication is inhibited by high Cdk levels in metaphase. Although high Cdk levels are compatible with bulk DNA replication in fission yeast (*2*), chromatin replication is inhibited in *Xenopus* mitotic extracts (*15*). Perhaps the replication of specific genomic regions must be paused during metaphase to prevent damage and chromosome pulverization (*16, 17*). Our data indicate that regions under-replicated in metaphase have one last opportunity to complete their replication before cytokinesis when Cdk levels fall below a critical threshold during mitotic exit.

Why do yeast not complete replication prior to mitosis? Starting mitosis before completion of DNA replication may allow for an increased cell division rate which could outweigh occasional mutations, aneuploidy and gene loss. For instance, a decrease in the number of viable cells can be easily compensated for by an accelerated division rate **(Fig. S14)**. In addition, our data raise the possibility that regions that replicate during mitosis, devoid of essential genes, serve as a genetic playground in which high rates of mutation and copy-number variation (*21*) allow cells to more rapidly explore genotypic space. Further, continued replication during late mitosis may explain the high frequency in animal cells of ultrafine chromatin bridges, whose defective resolution is associated with genome instability (*22*). Finally, our findings may have implications for how animal cells with rapid divisions such as in the early embryo, with no gap phases, ensure complete DNA replication.

## Acknowledgements

We’d like to thank the UPF/CRG Flow Cytometry core, the CRG sequencing and microscopy core facilities, and Mercè Gomar-Alba for assistance with microscopy; Yves Barral, Damien Coudreuse, Bruce Futcher, Travis Stracker, Jordi Torres-Rosell, Jenny Wu and Philip Zegerman for helpful comments; and Life Science Editors for editorial assistance. This study was supported by Ministerio de Economía y Competitividad (MINECO) (BFU2015-68351-P) and AGAUR (2014SGR0974 & 2017SGR1054) grants to L.B.C. and the Unidad de Excelencia María de Maeztu, funded by the MINECO (MDM-2014-0370); the European Research Council (ERC) Starting Grant 2010-St-20091118 to Ma.M., the Spanish Ministry of Economy and Competitiveness, ‘Centro de Excelencia Severo Ochoa 2013–2017’, SEV-2012-0208 to the CRG and the grant ANR-10-LABX-0030-INRT, which is a French State fund managed by the Agence Nationale de la Recherche under the frame programme Investissements d’Avenir ANR-10-IDEX-0002-02 to the IGBMC. M.M. was supported by a “La Caixa” PhD fellowship.

## Author Contributions

Ma.M. conceived the initial project. T.I., Mi.M. and A.M. performed experiments. T.I., Mi.M., A.M., L.B.C. and Ma.M. analyzed the data. L.B.C. and Ma.M. acquired funding, supervised the study and wrote the manuscript with input from all authors.

## Data and materials availability

All strains and plasmids are available upon request. Raw and processed sequencing data are at NCBI GEO (GSE117268 Data and code used in this work are available at https://github.com/amissarova/Mendoza__ReplicationEvolution.

## Materials and Methods

### Strains and cell growth

*S*. *cerevisiae* strains are derivatives of S288c except when noted (**Supplementary Table S4**). Gene deletions and insertions were generated by standard one-step PCR-based Methods (*25*). *cdc28-as1* strains (Cdc28-F88G) were made using plasmid pJAU01-cdc28-as1 (a kind gift from Ethel Queralt) digested with Afl2 or EcoN1 for integration at the *CDC28* locus; *URA3*+ clones were selected and then *ura3-* pop-outs isolated in 5-FOA plates; final clones were confirmed by Sanger sequencing. *MET3pr-CDC20* strains were made with YiP22-MET3pr-*CDC20*-LEU2 (Ethel Queralt) digested with MscI for integration in the *CDC20* locus. *GALpr-CLB2db* (in which the first 240 nt of *CLB2* encoding the destruction box are truncated) was constructed in BY4741 by inserting the NATMX-GAL1pr cassette at the native *CLB2* locus. For arrest in G1, cells were grown in YPDA (yeast extract, peptone, dextrose and adenine) medium to log phase, synchronized with 20 μg ml–1 alpha-factor (Sigma-Aldrich) for 2 h. For release from the G1 block, cells were washed in fresh YPDA medium at the indicated temperature. For arrest in metaphase, *MET3pr-CDC20* cells were grown in SD+Glu (synthetic defined + glucose) medium lacking methionine until mid-log phase, and the media were supplemented with 2 mM methionine to repress *CDC20* for 3 h 30 min. For release, cells were washed twice and resuspended in minimal medium lacking methionine to induce *CDC20*. Alternatively, cells were incubated with 60 μg/ml nocodazole for 3 hours. For Cdc15 or Cdk inhibition, *cdc15-as1* or *cdc28-as1* cells were incubated with NA-PP1 2.5 μM. For temperature shifts, cells were grown and arrested at 25 °C, shifted to 37 °C 15 minutes before washes, and released in prewarmed medium at 37.5 °C. Otherwise cells were arrested and released at 30 °C. For induction of the *GAL1,10* promoter, cells were grown overnight in YPR (yeast extract, peptone, raffinose) medium to mid-log phase, and galactose was added to 0.5%. To determine DNA content with Sytox-Green, we used the staining protocol from (*26*).

### EdU incorporation in cells arrested in metaphase

EdU click and DAPI staining were carried out as in (*27*) with minor modifications. Briefly, yeast cells (W303 *RAD5 bar1 MET3pr-CDC20 GPDpr-TK(5x) ADH1pr-hENT1*) expressing thymidine kinase (TK) and equilibrative nucleoside transporter (hENT1) to allow for EdU incorporation, were arrested in metaphase by Cdc20 depletion, as indicated. Arrested cells were then pre-incubated with 25 μM 5-ethynyl-2’-deoxyuridine (EdU) and α-Factor (20 μg/ml) for 15 minutes. Half the culture was kept in metaphase (+Met) and the other washed and resuspended into minimal medium (– Met) supplemented with 25 μM EdU and with 20 μg/ml α-Factor to block cells in the next G1 phase. After one hour cells were fixed with 2% paraformaldehyde and processed for Sytox Green staining (for flow cytometry) and for DAPI staining and EdU labeling with Alexa Fluor 647 (for microscopy) as described (*27*). Cells were imaged in a Leica TCS SP5 confocal microscope (63x objective, 3 planes spaced 0.67 μm). Nuclear EdU incorporation was measured in maximum projections as the background-subtracted mean intensity in the nucleus (defined by DAPI staining). The background was measured in multiple small cytoplasmic areas to avoid signal from mitochondrial DNA. As not all cells are permeabilized during fixation, we discarded those cells that showed no EdU signal in mitochondria.

### EdU incorporation in freely-cycling cells

was carried out in the same manner as above, in cells with wild-type CDC20 (w303, *RAD5 bar1 GPDpr-TK(5x) ADH1pr-hENT1*), except that cells were pregrown in SCD to mid-log phase and α-Factor (20 μg/ml) was added for 30 minutes to prevent cells already in G1 from entering into S phase. Cells were then incubated in EdU (25 μM) for 10 minutes; EdU was visualized with Alexa 647 and nuclei were visualised with DAPI (*27*).

### Microscopy

For time-lapse imaging, cells were plated in minimal synthetic medium on concanavalin A–coated (Sigma-Aldrich) Lab-Tek chambers (Thermo Fisher Scientific). Imaging was performed in a pre-equilibrated temperature-controlled microscopy chamber, using a spinning-disk confocal microscope (Revolution XD; Andor Technology) with a Plan Apochromat 100×, 1.45 NA objective equipped with a dual-mode electron-modifying charge-coupled device camera (iXon 897 E; Andor Technology). Time-lapse series of 4.5 μm stacks spaced 0.3 μm were acquired every 2 to 8 minutes. The time interval did not affect nuclear division or chromatin bridge lifetimes. iQ Live Cell Imaging software (Andor Technology) was used for image acquisition. Images were analyzed on 2D maximum projections. Maximum projections are shown throughout. For YOYO-1 quantification of chromatin bridges, cells were fixed in 4% formaldehyde for 30 min, washed once in PBS and re-suspended in 5mg/ml zymolyase in P solution (1.2M Sorbitol, 0.1M potassium phosphate buffer pH6.2) for 1 minute. Cells were spun down, taken up in P-Solution + 0.2% Tween 20 + 100 ug/ml RNAse A and incubated for 1h at 37 °C. After digestion, cells were pelleted and taken up in P-Solution containing 25 μM YOYO-1. Cells were imaged on an Andor Spinning disk microscope (15 planes spaced 0.3 μm).

## Quantification of under-replication across the genome by genome sequencing

To identify genomic regions that are under-replicated in mitosis, we performed whole-genome Illumina sequencing of cells arrested in metaphase (MET3pr-*CDC20*) or anaphase (*dbf2-2*). Each strain was also arrested in G1 phase (alpha factor arrest). *MET3pr-CDC20* cells were grown to mid-log phase in SC-Met, arrested in G1 with alpha factor at 25 °C for 3 hours, then washed and shifted to YPDA+Met for 3 hours at 37 °C to arrest in metaphase. *dbf2-2* cells were grown in YPDA, arrested in G1 with alpha factor at 25 °C for 3 hours, then washed into YPDA at 37°C to arrest in anaphase. For data shown in Figure 4A-E, cells were harvested by centrifugation for 5 min, washed twice in PBS, resuspended in water and layered on a 10%-40% sucrose gradient. After centrifugation at 400 g for 4 minutes, the top layers were removed, the remaining 10 ml of the gradient were split into two fractions and the pellet was taken up in 1 ml water to form the third fraction. Aliquots were removed from all three fractions to assay for cell cycle synchrony and the rest was pelleted and frozen at −80°C. Cell purification steps were performed at 37 °C. The fraction with the highest synchrony was used for phenol-chloroform-isoamyl DNA extraction. For data shown in figure 4F, cells were treated as above, except that Cdc20-depleted cells were shifted to 37°C during 15 minutes to inactivate *CDC12*, and then incubated with 1-NAA-PP1 (1 μM) for a further 30 minutes before DNA extraction with a gDNA extraction kit (Zymo Research). Cell synchronisation was assessed by flow cytometry and by microscopy (DAPI staining).

DNA sequencing was performed using Illumina HiSeq Sequencing V4 Chemistry with paired-end 50-bp reads. Reads were trimmed using Trimmomatic default parameters (java -Xms3G -Xmx3G -jar trimmomatic.jar PE -phred33) (*28*). Mapping was performed onto the R64 reference genome (*29*) of *S. cerevisiae* using Burrows-Wheeler Aligner with default parameters (bwa mem) (*30*). Reads were then sorted and improperly paired reads discarded (samtools view -F 4) using Samtools (*31*). We generated 200-bp non-overlapping windows across the genome (bedtools makewindows) and computed coverage for each window (bedtools coverage). Detailed and documented pipeline can be found in the Makefile on github and in the Supplementary Info.

Normalized coverage for each window was computed separately for each sequenced sample and each chromosome by dividing the read coverage for each window on that chromosome by the mode of all coverage values for all windows on that chromosome (copy number or CN). To each CN we applied a local regression using a weighted linear least squares and a first degree polynomial model (with a 5 kb span) to decrease the impact of technical noise. For each mutant we calculated copy number ratio (CNR) for each 200-bp window as normalized copy number in M-phase divided by the normalized copy number in G1-phase. The percent of cells that have not yet replicated a given region of the genome (under-replication), is defined as 2*(1-CNR).

The above was performed for six biological replicates for each strain and condition (see Supplementary Table S1), one outlier replicate was removed, and the median value of under-replication across replicates was assigned as a final value for each 200-bp window. All raw FASTQ files together with smoothed CN and datasets with calculated statistics for each 200-bp genomic window are on github and NCBI GEO.

## Analysis of under-replicated regions

To define which genomic regions (at the resolution of 200-bp) were significantly under-replicated we compared the actual CNRs between M-phase and G1-phase against technical noise. To generate a technical and biological noise CNR dataset we took CN data from pairs of G1 arrested biological replicates and calculated CNR (CNR-G1-control) between them in the same manner we did for M-and G1-phase (overall 15 pairs, for each pair - 2 CNR vectors: CN(replicate #1)/CN(replicate #2) and CN(replicate #2)/CN(replicate #1)). Then for each 200-bp window, starting from the first 200-bp from each telomeric end, we compared values of actual CNR vectors (5 in total, one for each repeatable replicate) against CNR-G1-control vectors at this window (Student T-test). The first window, starting from the telomeric end, where difference was not significant (p-value < 0.05) or the median for CN-ratios between samples in G1 was higher than median of actual CNR-value, was marked as the end of under-replicated region.

To choose a threshold value of under-replication at which regions of the genome are under-replicated in metaphase we used the under-replication distribution of all 200-bp windows. This distribution has a heavy right tail and positive skewness. The threshold value selected to classify loci as being under-replicated is the highest under-replication value where skewness of values lower than the selected value is 0.

## Quantification of under-replication for genomic features

To extract a single value for different genomic features of varying length, we first extracted coordinates of the features of interest. We compiled a dataset from different sources, including SGD (https://www.yeastgenome.org for centromeres tRNAs, transposons and ORFs in general) and OriDB (http://cerevisiae.oridb.org) for ARSs. ARSs marked as dubious were excluded from the analysis. Additionally, we added regions that were likely to undergo loss of heterozygosity (LOH) as well as duplications and deletions during cell growth (*14*). We denoted as fragile sites regions marked as either interstitial or terminal deletions or duplications in (*14*). For all collected features we extracted a single value of under-replication from the 200-bp resolution CNR vector using bedtools software (bedtools coverage). Replication timing data were extracted from (*23*) and spline interpolation was used to define a single replication timing value for each 200-bp window. To predict the occurrence of G-quadruplexes throughout the genome we used G4-Hunter (*32*) with a window size of 25nt (the default) and a score threshold of 1 (python G4Hunter.py -i yeastgenome.fasta –o S1_W25 -w 25 -s 1). For each 200 nt window we took the sum of all sequences that passed the score threshold (bedtools map -c 5 -o sum -a genome_windows_by_200.bed -b <G4HunterOutputFile.nts>). To calculate average under-replication and distance to telomeric start on gene ontology level, we extracted database with records of all ontologies and corresponding ORFs (not nested) from SGD, and for each ontology we calculated mean and CV (std/mean) for under-replication and distance to telomeric start for all genes in each GO term. The relative fitness for all essential genes was assigned as 0 while the measured fitness for non-essential genes are from (*33*). To extract representative metrics of genetic divergence between strains (mutations and copy-number variation) we used whole-genome sequences of 85 strains from (*24*). For each 200 bp-window we calculated frequency of SNPs and Indels. The frequency is the # of positions in each 200 nt window that have a SNP or Indel in at least one strain. Additionally, for each gene, we calculated a fraction of strains in which this gene is present at the given locus.

